# A novel temporal identity window generates alternating cardinal motor neuron subtypes in a single progenitor lineage

**DOI:** 10.1101/2020.02.12.946442

**Authors:** Austin Seroka, Rita M Yazejian, Sen-Lin Lai, Chris Q Doe

## Abstract

Spatial patterning specifies neural progenitor identity, with further diversity generated by temporal patterning within individual progenitor lineages. These mechanisms generate cardinal classes of motor neurons (sharing a transcription factor identity and common muscle group targets). In *Drosophila*, two cardinal classes are Even-skipped (Eve)+ motor neurons projecting to dorsal muscles and Nkx6+ motor neurons projecting to ventral muscles. The *Drosophila* neuroblast 7-1 (NB7-1) lineage generates distinct Eve+ motor neurons via the temporal transcription factor (TTF) cascade Hunchback (Hb)-Krüppel (Kr)-Pdm-Castor (Cas). Here we show that a newly discovered Kr/Pdm temporal identity window gives rise to an Nkx6+ Eve-motor neuron projecting to ventral oblique muscles, resulting in alternation of cardinal motor neuron subtypes from a single progenitor (Eve>Nkx6>Eve). We show that co-overexpression of Kr/Pdm generates ectopic VO motor neurons within the NB7-1 lineage and that Kr/Pdm act via Nkx6, which itself is necessary and sufficient for VO motor neuron identity. Lastly, Nkx6 is required for ventral oblique muscle targeting, thereby linking temporal patterning to motor neuron morphology and synaptic target selection. In conclusion, we show that one neuroblast lineage generates interleaved cardinal motor neurons fates; that the Kr/Pdm TTFs form a novel temporal identity window that promotes expression of Nkx6; and that the Kr/Pdm>Nkx6 pathway is necessary and sufficient to specify VO motor neuron identity and morphology.

## Introduction

Neural diversity from flies to mice arises from two major developmental mechanisms. First, neural progenitors acquire a unique and heritable spatial identity based on their position along the rostrocaudal or dorsoventral body axes (Kohwi and Doe, 2013; Sagner and Briscoe, 2019). Second, temporal patterning based on neuronal birth-order results in individual progenitors producing a diverse array of neurons and glia (Holguera and Desplan, 2018; Kohwi and Doe, 2013; Miyares and Lee, 2019). Temporal patterning is best characterized in *Drosophila*; neural progenitors (neuroblasts) located in the ventral nerve cord, central brain, and optic lobes all undergo temporal patterning, in which the neuroblast sequentially expresses a cascade of TTFs that specify distinct neuronal identities (Allan and Thor, 2015; Doe, 2017; Holguera and Desplan, 2018; Miyares and Lee, 2019). Although all neuroblasts undergo temporal patterning, the TTFs are different in each region of the brain (Allan and Thor, 2015; Doe, 2017; Holguera and Desplan, 2018; Miyares and Lee, 2019). Similar mechanisms are used in the mammalian cortex, retina, and spinal cord, although many TTFs remain to be identified (Alsiö et al., 2013; Delile et al., 2019; Elliott et al., 2008; Mattar et al., 2015; Sockanathan and Jessell, 1998; Stam et al., 2011).

A major open question is how transient expression of TTFs like Kr and Pdm lead to long-lasting specification of molecular and morphological neuronal diversity. It is likely that TTFs induce expression of suites of transcription factors that persist in neurons and confer their identity, but in only a few cases have these “morphology transcription factors” been identified, such as those specifying adult leg motor neuron dendrite projections (Enriquez et al., 2015), but in this case little is known about the earlier TTFs that establish morphology transcription factor expression. Conversely, the TTF cascade is well established in the ventral nerve cord neuroblasts, but less is known about downstream transcription factors that consolidate neuronal identity. Good candidates are the transcription factors known to specify cardinal motor neuron identity. The Eve+ cardinal motor neurons project to dorsal muscles, while the cardinal motor neurons expressing Nkx6, Hb9, Islet, or Lim3 target ventral muscles (Broihier and Skeath, 2002; Broihier et al., 2004; Certel and Thor, 2004; Landgraf et al., 1999; Venkatasubramanian and Mann, 2019). Dorsal and ventral motor neuron cardinal classes exhibit cross-repression to consolidate their dorsal or ventral identity (Broihier and Skeath, 2002).

The *Drosophila* neuroblast 7-1 (NB7-1) is arguably the best characterized system for understanding TTF expression and function. Similar to most other ventral nerve cord neuroblasts, NB7-1 expresses the canonical TTF cascade Hb-Kr-Pdm-Cas with each TTF inherited by the GMCs born during an expression window, and transiently maintained in the two post-mitotic neurons produced by each GMC. The TTF cascade generates diversity among the five Eve+ U1-U5 motor neuron progeny of NB7-1: Hb specifies U1 and U2, Kr specifies U3, Pdm specifies U4, and Pdm/Cas together specify U5 (Cleary and Doe, 2006; Grosskortenhaus et al., 2006; Isshiki et al., 2001; Pearson and Doe, 2003). Identifying TTF target genes, including transcription factors and cell surface molecules, will provide a comprehensive view of how developmental determinants direct neuronal morphology and synaptic partner choices.

It has long been thought that the dorsal and ventral cardinal classes of motor neurons derive from distinct progenitors; Eve+ motor neurons derive from NB7-1, NB1-1, and NB4-2 whereas Hb9+ or Nkx6+ motor neurons derive from NB3-1 and others. However, DiI labeling of NB7-1 identified an unknown motor neuron innervating ventral muscles, which is distinct from dorsal muscles targeted by the Eve+ motor neurons (Schmid et al., 1999). Ventral muscle innervation could be transient exuberant outgrowths that are subsequently pruned, or uncharacterized motor neurons that form stable synapses with ventral muscles.

Here, we show that a newly discovered Kr/Pdm TTF window generates an Nkx6+ Eve-ventral projecting VO motor neuron, born between U3 and U4 in the NB7-1 lineage. We also show that overexpression of Kr/Pdm together, or Nkx6 alone, generates ectopic ventral projecting VO motor neurons. Finally, we demonstrate that Nkx6 is required for proper VO motor neuron axon targeting to ventral oblique muscles. Our results establish a genetic pathway from TTFs (Kr/Pdm), to a cardinal motor neuron transcription factor (Nkx6) to axonal targeting. This is among the first TTF target genes known to regulate neuronal morphology and connectivity. We also make the unexpected discovery that a single progenitor can alternate production of different cardinal motor neuron classes.

## Results

### The NB7-1 lineage has a Kr+ Pdm+ temporal identity window that generates an Nkx6+ motor neuron

To determine the identity of the neurons originating from the newly identified Kr+ Pdm+ temporal identity window, we used a previously-characterized highly-specific NB7-1 split gal4 line (*NB7-1-gal4*; Seroka and Doe, 2019) to express *UAS-GFP* in NB7-1 and its progeny (Figure 1A-D). This driver line is expressed during the early part of the lineage, including the time U1-U5 neurons are born, but fades out before the end of the lineage (Seroka and Doe, 2019). We used Eve to identify the U1-U5 motor neurons within the lineage, and Zfh1 to label all motor neurons (Layden et al., 2006). We identified a single Eve-Zfh1+ motor neuron in the lineage (Figure 1A-D). This Eve-Zfh1+ “sixth motor neuron” was Kr+Pdm+ (Figure 1A,B), consistent with originating from the previously defined Kr+Pdm+ GMC within the NB7-1 lineage (Averbukh et al., 2018). To determine whether the Kr+Pdm+ motor neuron is a U1-U5 sibling, we stained for the Notch reporter gene Hey. All U1-U5 motor neurons are Hey+ and have Hey-sibling neurons (Skeath and Doe, 1998). We confirmed that all U1-U5 motor neurons express the Notch reporter Hey, and show that the Kr+Pdm+ motor neuron is also Hey+ (Figure 1C). Thus, the Kr+Pdm+ motor neuron and Eve+ U1-5 motor neurons are each derived from a distinct GMC in the lineage, consistent with the Kr+Pdm+ motor neuron originating from the fourth-born Kr+Pdm+ GMC (Averbukh et al., 2018).

**Figure 1.**
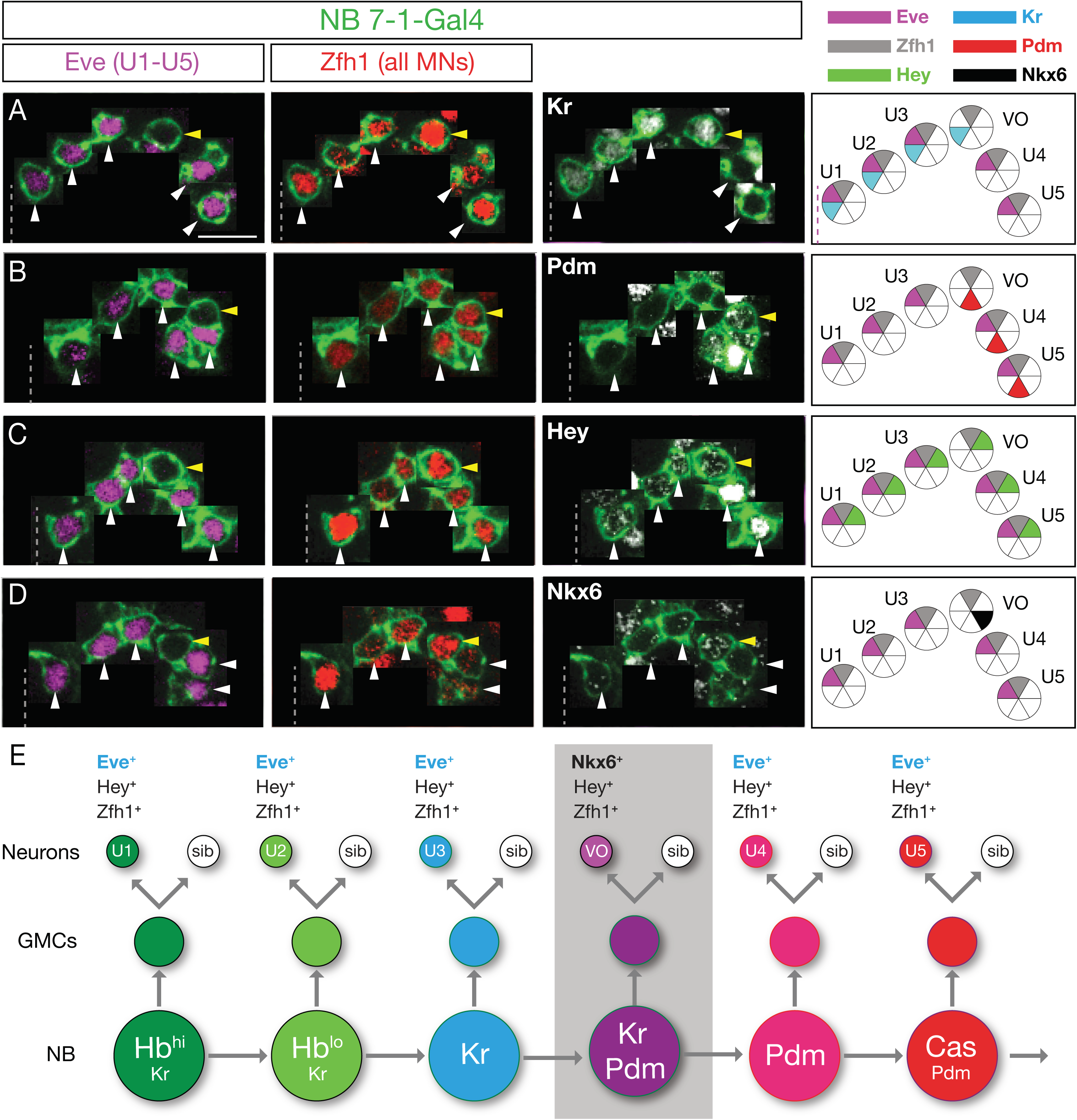
NB7-1 generates an Eve- Nkx6+ motor neuron. In this and all following panels, one hemisegment of a stage 16 embryo; ventral midline, dashed line; Eve+ Nkx6- motor neurons, white arrowhead; Nkx6+ Eve- motor neuron, yellow arrowhead. Neurons are montaged from different z-axis positions with their X-Y position preserved (see Methods) (A-D) The NB7-1 lineage is labeled with GFP in all columns (*NB7-1-gal4 UAS-GFP*; green). Left column: Eve identifies U1-U5 motor neurons. Middle column: Zfh1 identifies all motor neurons, although in U2 motor neurons, Zfh1 staining is fainter. Right column shows the indicated marker. Far right column: summary. All images from stage 16/17 abdominal segments. For Nkx6 stain all 6 cells are Hey+, however the intensity in much higher in the U4/U5 motor neurons born most recently at stage 17. Scale bar: 10μm. (E) Proposed NB7-1 lineage.

The transcription factors Eve and Nkx6 have cross-repressive interactions (Broihier et al., 2004), raising the possibility that the Kr+Pdm+ motor neuron might be Nkx6+. Indeed, we confirmed that the Kr+Pdm+ motor neuron is Nkx6+ (Figure 1D). Interestingly, this Nkx6+ VO motor neuron is negative for other ventral neuron markers, including Hb9, Islet and Lim3 (data not shown). We conclude that the NB7-1 lineage produces two cardinal classes of motor neurons: dorsal projecting Eve+ motor neurons and ventral projecting Nkx6+ motor neurons; unexpectedly, these cardinal classes of motor neurons are produced in an alternating mode from a single progenitor lineage: 3 Eve motor neurons > 1 Nkx6 motor neuron > 2 Eve motor neurons (Figure 1E). This is surprising, and one of the few examples of a progenitor alternating cell types within its lineage (see Discussion).

### The Nkx6+ motor neuron projects to ventral oblique muscles

All of the Nkx6+ motor neurons that have been characterized to date project to ventral body wall muscles (Broihier et al., 2004). To identify the muscle target of the Kr+Pdm+Nkx6+ motor neuron in the NB7-1 lineage, we first examined single neuroblast DiI clones (Schmid et al., 1999) and looked for muscles with innervation distinct from the known U1-U5 dorsal muscle targets. DiI labeling of NB7-1 marked all clonal progeny at embryonic stage 17, and showed innervation of all known U1-U5 dorsal muscle targets, plus innervation of the ventral oblique muscles 15, 16, and 17 via the intersegmental nerve d branch (ISNd) (Figure 2A). We independently confirmed these results using NB7-1-gal4 to drive GFP expression, which labeled all U1-U5 dorsal muscle targets plus ventral oblique muscles in a majority of hemisegments (Figure 2B, C; 54/93 hemisegments). These data are consistent with the Nkx6+Eve-motor neuron in the NB7-1 lineage projecting to ventral oblique muscles, but without single neuron labeling we can’t make a conclusive match.

**Figure 2.**
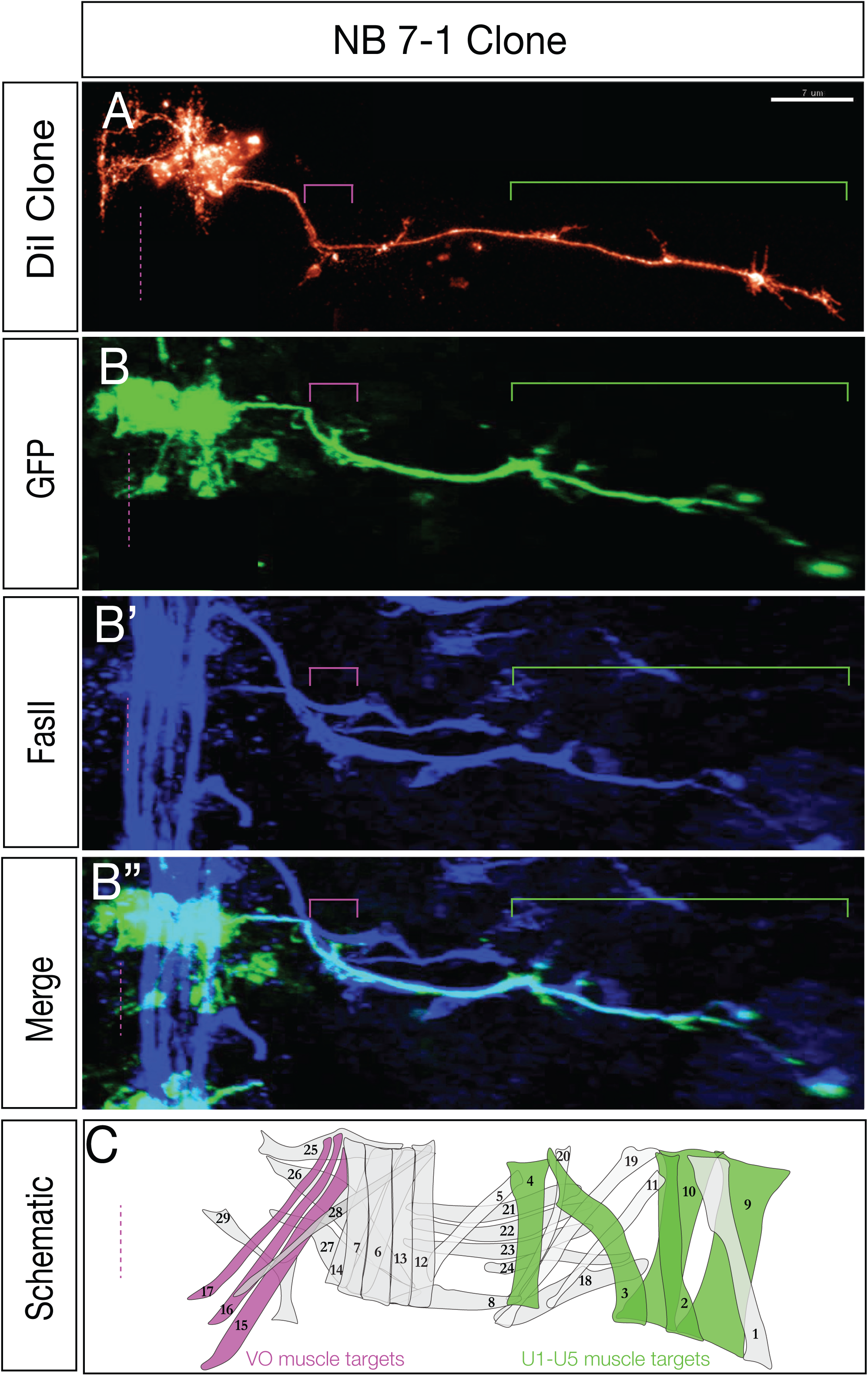
NB7-1 generates a motor neuron that innervates ventral oblique muscles. (A) NB7-1 DiI clone (red) generated as described in Schmid et al. (1999). Green bracket, U1-U5 motor neurons innervating dorsal muscles; magenta bracket, unknown neuron(s) innervating ventral muscles (not shown). (B-B”) NB7-1 lineage marked by GFP (*NB7-1-gal4 UAS-GFP*; green) and stained for the motor axon marker FasII, showing motor neuron innervation of dorsal (green bracket) and ventral (magenta bracket) muscle groups. Stage 17; ventral midline, dashed; dorsal body wall to right. Scale bar: 7 μm. (C) Schematic of dorsal and ventral muscle groups, including the ventral oblique muscles (magenta) and the dorsal longitudinal muscles (green).

To visualize the morphology of individual motor neurons in the NB7-1 lineage, we used multicolor flipout (MCFO) (Nern et al., 2015) within the NB7-1 lineage (see methods for full genotype). We used *NB7-1-gal4 UAS-GFP* to label all neurons in the lineage, FasII to detect all motor axons and their muscle targets, and individual HA+ neurons using MCFO (Figure 3A-A’’’). As expected, full lineage labeling revealed innervation of ventral oblique muscles via ISNd. In addition, HA labeling identified individual motor neurons that specifically targeted the ventral oblique muscles via ISNd. This motor neuron had an ipsilateral dendritic process that approached the midline (Figure 3A’), indistinguishable from the previously identified MN17 that innervates the ventral oblique muscles (Landgraf et al., 2003). Thus, we call the Kr+Pdm+Nkx6+ motor neuron in the NB7-1 lineage the “VO motor neuron” henceforth.

**Figure 3.**
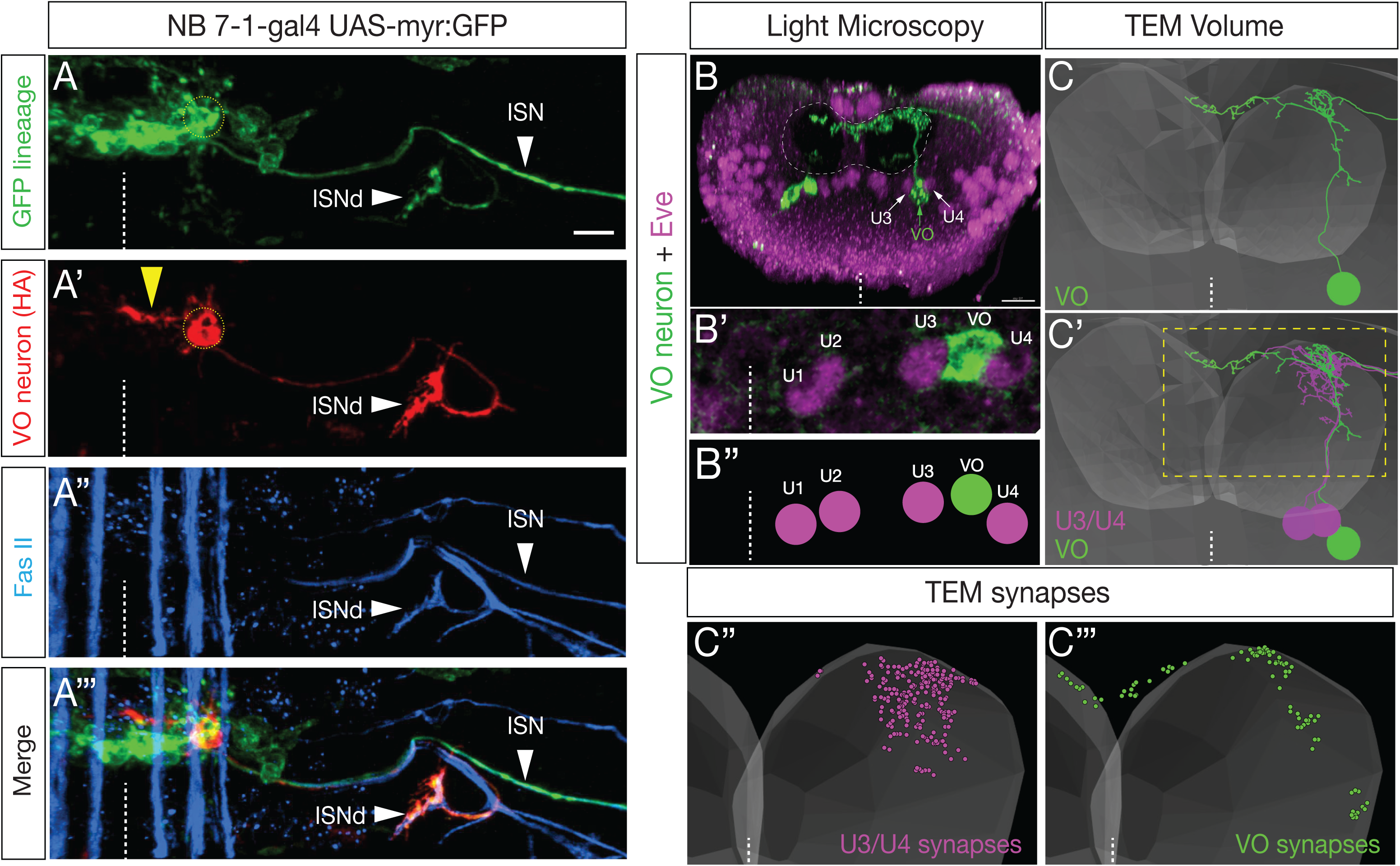
NB7-1 derived Nkx6+ motor neuron projects axon to ventral oblique muscles and dendrites to the dorsal neuropil. (A-A’’’) NB7-1 lineage marked by GFP (*NB7-1-gal4 UAS-GFP*; green) as well as a single neuron marked by MCFO (HA; red) showing innervation of ISNd nerve root. FasII (blue) marks all motor projections. Stage 16 embryo; ventral midline, dashed line. Scale bar: 5 μm. (B-B’’) NB7-1 lineage MCFO showing a single VO motor neuron (green) in a newly hatched larva; posterior (cross-section) view; dorsal up, ventral midline, dashed line; neuropil boundary, dashed circle. VO motor neuron cell body (green arrow) lies adjacent to U3 and U4 neuron cell bodies (white arrows). Note the VO motor neuron dendrite at the dorsal-most neuropil. Scale bar: 10 μm. (C-C’’’) VO motor neuron (green) is identifiable in a TEM reconstruction of the newly hatched larval CNS, where it was called MN15/16/17 (Zarin et al., 2019). (C’) Note the similar morphology and cell body position compared to VO in panel B. Posterior (cross-section) view; dorsal up, ventral midline, dashed line; approximate neuropil boundary, translucent shading. U3/U4 cell bodies (magenta) are adjacent to the VO cell body (green). (C’’, C’’’) Presynapse locations for premotor inputs to VO (green) and U3/U4 (magenta).

We next wanted to determine whether the Nkx6+ VO motor neuron and the Eve+ U1-U5 motor neurons have distinctive dendritic morphology or premotor innervation, as expected based on their distinctive axon targeting to different muscle groups. We repeated the MCFO experiment in newly hatched larvae so we could identify VO motor neuron morphology in a recently described TEM atlas of all abdominal motor neurons performed in newly hatched larvae (Zarin et al., 2019) (Figure 3A-B’’). We identified the NB7-1 VO motor neuron in the TEM volume based on its characteristic CNS projections and cell body position between U3 and U4 (compare Figure 3B and 3C). Interestingly, VO and U3/U4 motor neurons had different dendrite projections (Figure 3C), as well as different postsynapse locations (Figure 3C’’, C’’’). Furthermore, they had distinctive premotor inputs: the top three inputs to VO are A27h, A06c, A18b2 interneurons whereas the top three inputs to U3/U4 are A31k, A18a, and A31b interneurons (Zarin et al., 2019). We conclude that the NB7-1 lineage produces two cardinal classes of motor neurons: dorsal projecting Eve+ motor neurons and a ventral projecting Nkx6+ VO motor neuron, each with distinct morphology and connectivity.

### Overexpression of Kr/Pdm generates ectopic Nkx6+ VO motor neurons

Hb, Kr, Pdm, Pdm/Cas each can specify a distinct temporal identity within multiple neuroblast lineages (Doe, 2017). In contrast, the newly discovered Kr/Pdm temporal identity window (Averbukh et al., 2018) has not yet been tested for a role in specifying neuronal identity. Here we ask whether co-expression of Kr and Pdm can induce ectopic VO neurons. To test this hypothesis, we overexpressed Kr and Pdm together specifically in the NB7-1 lineage (*NB7-1-gal4 UAS-Kr UAS-Pdm UAS-GFP*). Controls always have a Nkx6+ Zfh1+ VO motor neuron located between U3 and U4 (Figure 4A; quantified in 4C). In contrast, Kr/Pdm co-expression resulted in ∼2 additional Nkx6+ Zfh1+ VO motor neurons (Figure 4B; quantified in 4C). We observed similar results with a second line, *engrailed (en)-gal4* (Figure 4G,H; quantified in 4I,J). We note that Kr/Pdm overexpression is not able to alter earlier temporal identities (Hb+ U1/U2 and Kr+ U3 neurons), similar to the well-characterized inability of later TTFs to alter earlier TTF cell fates (Cleary and Doe, 2006; Grosskortenhaus et al., 2006; Isshiki et al., 2001; Pearson and Doe, 2003; Tran and Doe, 2008). We conclude that Kr/Pdm TTFs are sufficient to induce Nkx6+ motor neuron identity beginning with the fourth division of the NB7-1 lineage.

**Figure 4.**
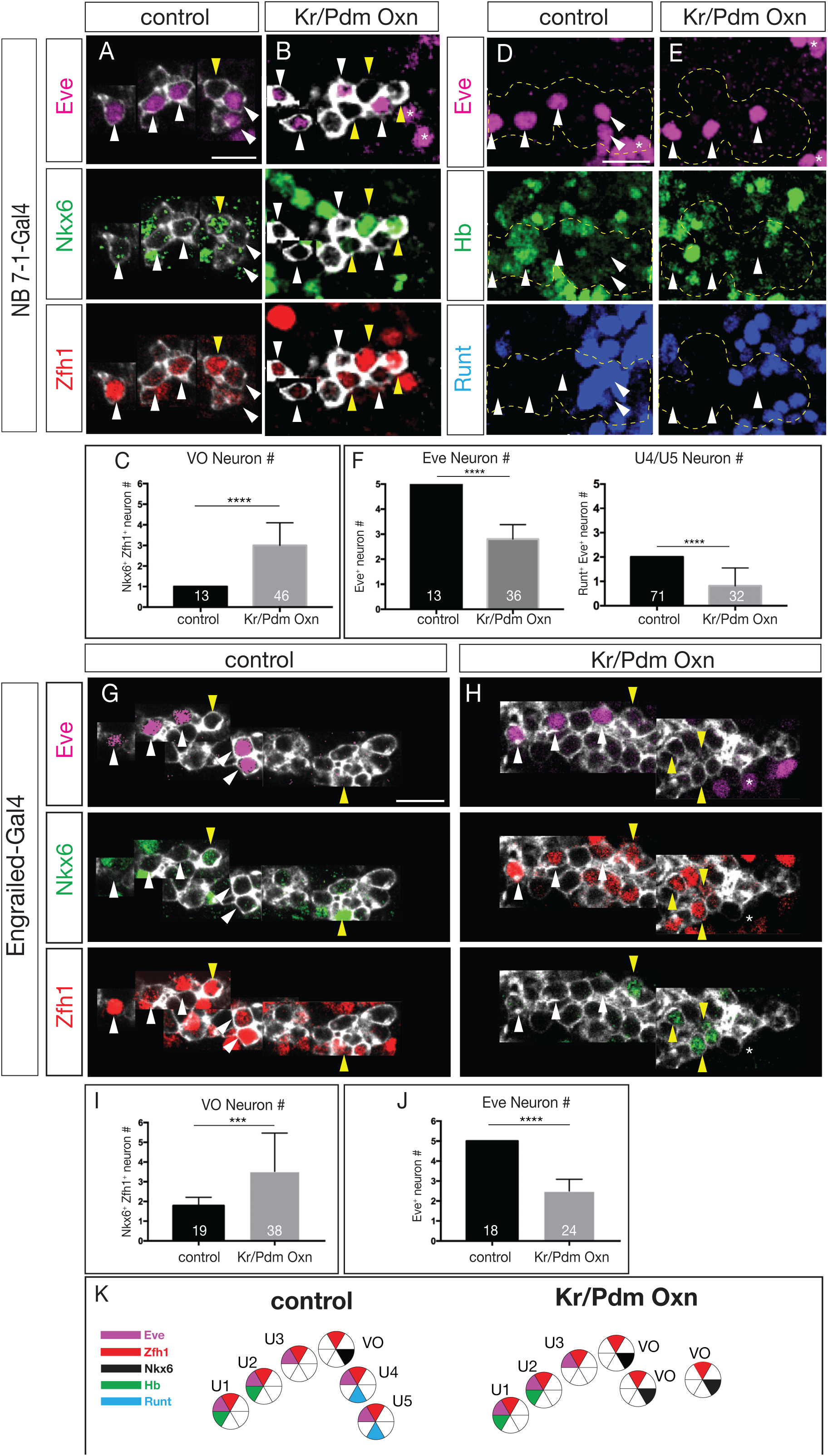
Kr/Pdm overexpression induces ectopic Nkx6+ VO motor neuron identity. In this and all following panels, one hemisegment of a stage 16 embryo; ventral midline, dashed line; VO motor neuron, yellow arrowhead; U motor neurons, white arrowhead; asterisk, EL neurons from NB3-3. Neurons are montaged from different z-axis positions with their X-Y position preserved (see Methods) (A-C) NB7-1-gal4 driving Kr/Pdm overexpression (Oxn) induces ectopic Nkx6+ VO motor neurons. (A) Control lineages have one Nkx6+ Eve- VO motor neuron. In this panel and below, neurons are montaged from different z-axis positions with their X-Y position preserved (see Methods). (B) Overexpression of Kr/Pdm together increases the number of Nkx6+ Eve- VO motor neurons. (C) Quantification. Scale bar: 10 μm. (D-F) NB7-1-gal4 driving Kr/Pdm overexpression (Oxn) induces loss of Eve+ motor neurons. Dashed outline, boundary of lineage marked with GFP (*NB7-1-gal4 UAS-GFP*). (D) Control lineages have five Eve+ motor neurons including Hb+ U1/U2 and Runt+ U4/U5. (E) Overexpression of Kr/Pdm together reduces the number of Eve+ U4/U5 motor neurons. (F) Quantification. Scale bar: 10 μm. (G-J) en-gal4 driving Kr/Pdm overexpression (Oxn) induces ectopic Nkx6+ VO motor neurons and loss of Eve+ motor neurons. (G) Controls have five Eve+ U1-U5 motor neurons and one Nkx6+ VO motor neuron. (H) Overexpression of Kr/Pdm together increases the number of Nkx6+ Eve- VO motor neurons. (I,J) Quantification. Scale bar: 10 μm. (K) Summary of Kr/Pdm overexpression phenotype.

We next asked whether Kr/Pdm co-expression delays production of the later-born Eve+ U4-U5 motor neurons until after birth of the ectopic VO motor neurons (lineage extension model), or replaces the Eve+ U4-U5 motor neurons with ectopic VO motor neurons (conversion model). Both outcomes have been observed following misexpression of other temporal transcription factors (Kohwi et al., 2013; Meng et al., 2019; Pearson and Doe, 2003; Seroka and Doe, 2019). We overexpressed Kr/Pdm in the NB7-1 lineage (*NB7-1-gal4 UAS-Kr UAS-Pdm UAS-GFP*), or more broadly (*en-gal4 UAS-Kr UAS-pdm*) and assayed the fate of the Eve+ U1-U5 motor neurons. In controls, we always observed five Eve^+^ U1-U5 motor neurons, including two U4/U5 motor neurons (Figure 4D,G; quantified in 4F,J). In contrast, Kr/Pdm overexpression led to a loss of the Runt+ U4/U5 motor neurons (Figure 4E,H; quantified in 4F,J). We conclude that overexpression of Kr/Pdm in the NB7-1 lineage generates ectopic Nkx6+ VO motor neurons at the expense of the later-born U4/U5 motor neurons, supporting the conversion model (summarized in Figure 4K).

### Overexpression of Kr/Pdm generates ectopic Nkx6+ motor neurons that project correctly to ventral oblique muscles

In the section above we show Kr/Pdm overexpression can induce ectopic VO motor neurons based on molecular markers. Here we determine whether Kr/Pdm overexpression can generate ectopic VO motor neurons that project correctly to ventral oblique muscles. We used *NB7-1-gal4* to drive prolonged overexpression of Kr/Pdm, membrane targeted GFP to visualize axon projections, and FasII to identify the ISNd branch to the ventral oblique muscles. In controls, NB7-1 progeny projected in the ISNd to innervate ventral oblique muscles (Figure 5A, S2A; axon volume in ISNd quantified in 5C; Figure S1). Following Kr/Pdm overexpression specifically in the NB7-1 lineage, we detected increased axon volume at the ISNd (Figure 5B, S2B; quantified in 5C), consistent with ectopic VO motor neurons taking the normal VO pathway via ISNd to the ventral oblique muscles. To conclusively show that multiple ectopic VO motor neurons target the ISNd, we used MCFO to express HA and V5 on different neurons within the NB7-1 lineage. Following Kr/Pdm overexpression specifically in the NB7-1 lineage, we identified NB7-1 lineages with distinct HA+ and V5+ motor neurons that projected via ISNd to the ventral oblique muscles (Figure 5D). Importantly, both HA+ and V5+ VO motor neurons showed specific targeting to the ventral oblique muscles. We conclude that overexpression of Kr/Pdm is sufficient to generate VO motor neurons that correctly project out of the ISNd nerve root to innervate ventral oblique muscles. Thus, Kr/Pdm induces both molecularly and morphologically normal VO motor neurons.

**Figure 5.**
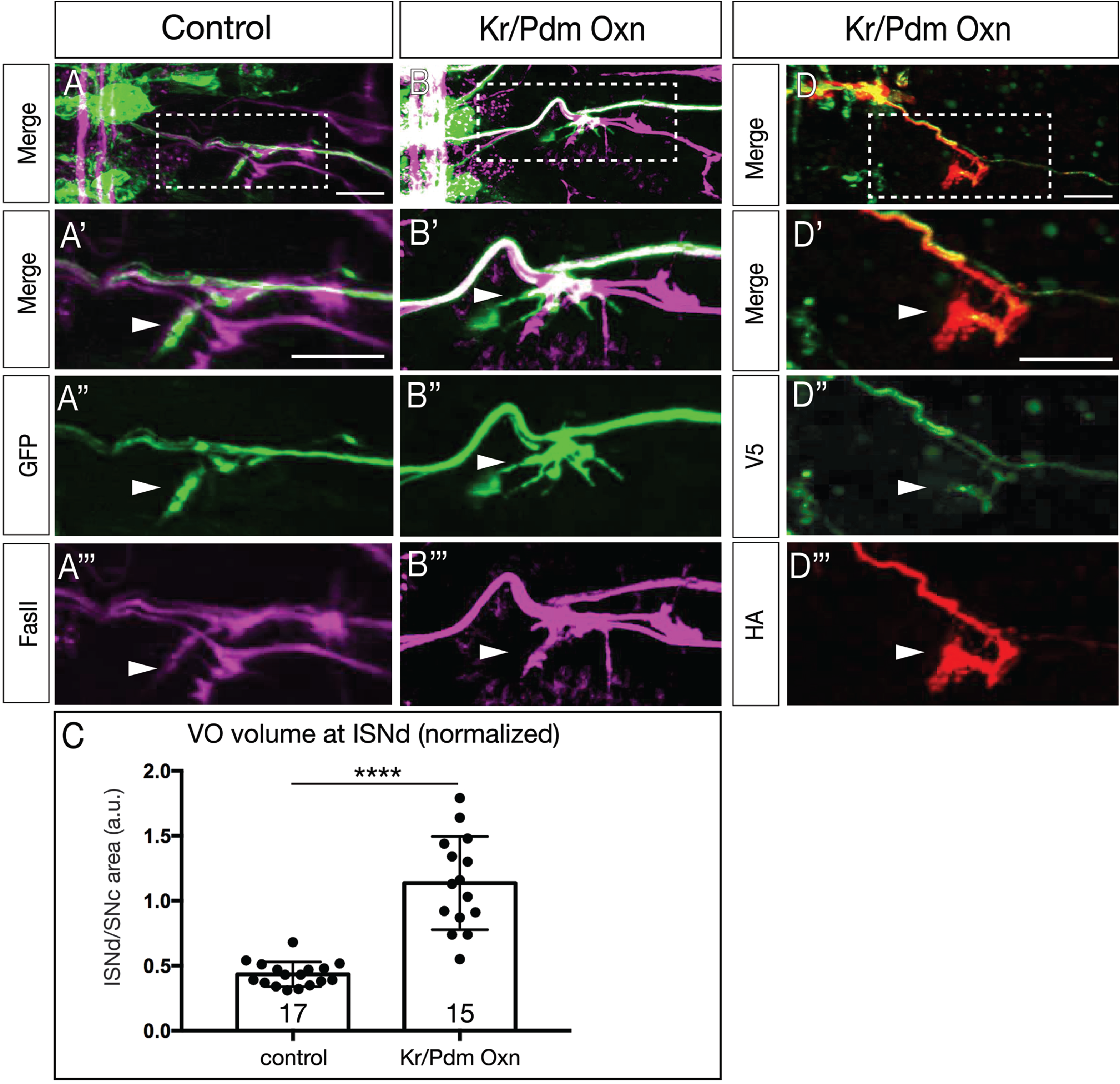
Kr/Pdm induces ectopic VO motor neurons targeting ventral oblique muscles. (A-C) Kr/Pdm overexpression results in ectopic motor neuron projections to the ventral oblique muscles (arrowhead). (A) Controls show NB7-1 progeny innervating the ventral oblique muscle (ISNd). (A’) Enlargement of boxed region in A. (A’’) GFP marking NB7-1 progeny; (A’’’) FasII marking all motor axons. (B) Overexpression of Kr/Pdm together (*NB7-1-gal4 UAS-Kr UAS-Pdm)* induces ectopic VO motor neuron projections to the ventral oblique muscles (arrow). Scale bar: 10 μm. (C) Quantification of ISNd volume normalized to SNc volume (ISNd/SNc; see Figure S1). (D) Kr/Pdm overexpression analyzed by single neuron MCFO labeling showing multiple VO motor neurons projecting to the ventral oblique muscles. (D’) Enlargement of boxed region in D. (D’’, D’’’) Individual motor neurons labeled with V5 or HA projecting to ventral oblique muscles. Scale bar: 10 μm.

### Nkx6 is necessary and sufficient to generate VO motor neurons in the NB7-1 lineage

We have shown above that the TTF combination of Kr/Pdm can induce Nkx6+ motor neurons within the NB7-1 lineage. This raises the question of whether there is a linear genetic pathway from Kr/Pdm to Nkx6 to VO motor neuron identity. This hypothesis is supported by previous work showing that Nkx6 overexpression can induce ectopic motor neurons with ventral muscle projections (Broihier et al., 2004; Cheesman et al., 2004). Thus, we asked if Nkx6 is necessary or sufficient to specify VO motor neuron identity. In wild type, *NB7-1-gal4* always detects the U1-U5 Eve+Zfh1+ motor neurons and a single Nkx6+Zfh1+ VO motor neuron (Figure 6A). When we use *NB7-1-gal4* to drive expression of UAS-Nkx6, we detect a loss of U3-U5 Eve+ motor neurons and the production of ectopic VO motor neurons based on molecular marker expression (Figure 6B; quantified in 6E). Conversely, when we use *NB7-1-gal4* to drive expression of *UAS-Nkx6-RNAi*, we detect a loss of the Nkx6+Zfh1+ VO motor neuron and the production of a single ectopic “U3.5” Eve+Zfh1+ motor neuron (Figure 6C; quantified in 6E). We conclude that Nkx6 is necessary and sufficient for specifying the molecular identity of the VO motor neuron in the NB7-1 lineage.

**Figure 6.**
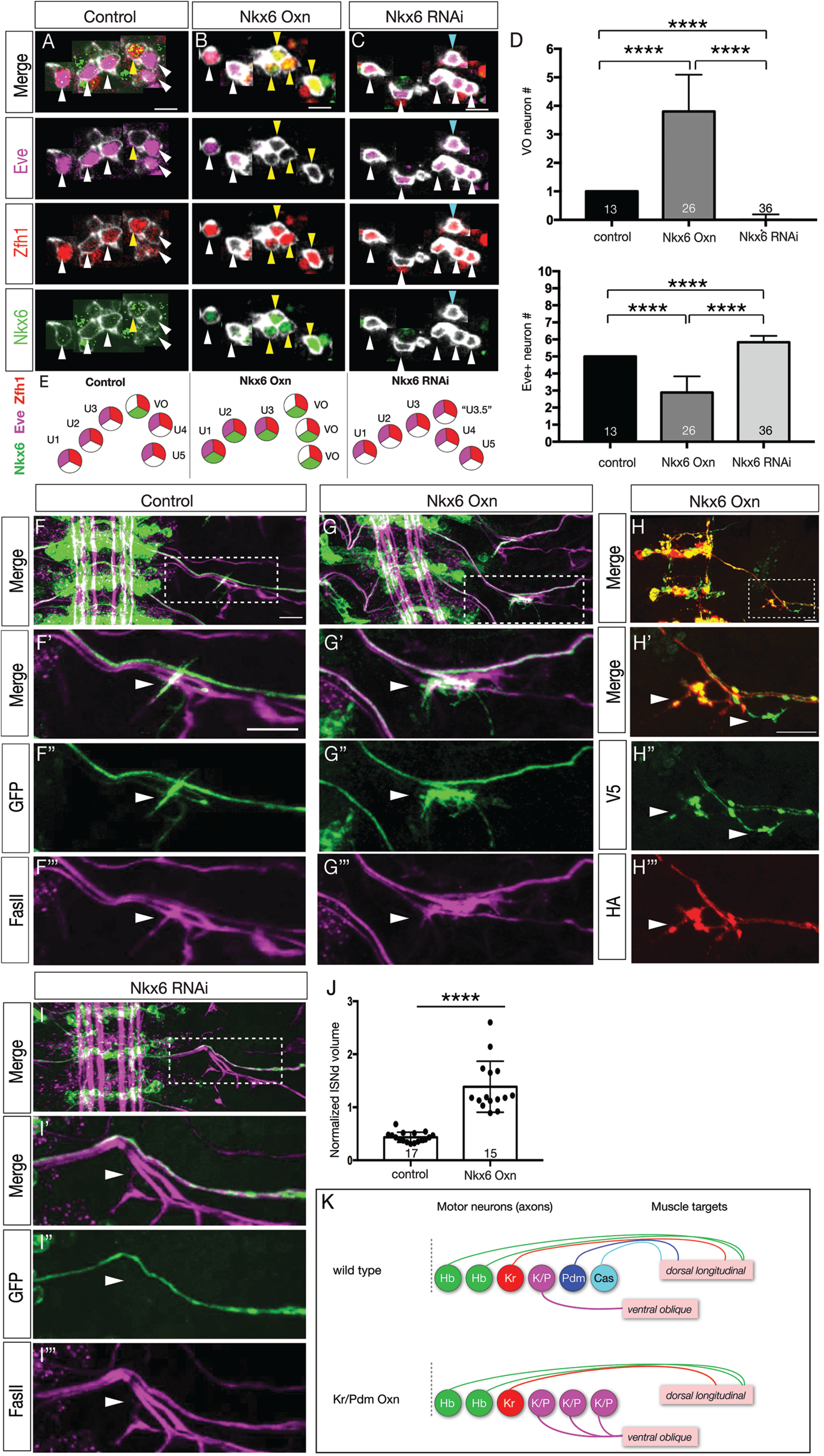
Nkx6 is necessary and sufficient to specify VO motor neuron identity. In this and following panels, one hemisegment of a stage 16 embryo; ventral midline, left side of panel; VO motor neuron, yellow arrowhead; U motor neurons, white arrowhead; ectopic Eve+ neuron at the normal VO location, cyan arrowhead. Neurons are montaged from different z-axis positions with their X-Y position preserved (see Methods) (A-E) Nkx6 specifies VO motor neuron molecular identity. (A) Controls have one Nkx6+ Eve- VO motor neuron; (B) Nkx6 overexpression in the NB7-1 lineage (*NB7-1-gal4 UAS-Nkx6*) have ectopic VO motor neurons at the expense of Eve+ U3-U5 motor neurons; (C) Nkx6 RNAi in the NB7-1 lineage (NB7-1-gal4 UAS-Nkx6-RNAi) lack the VO motor neuron and possess an ectopic Eve+ “U3.5” motor neuron at that location (cyan arrowhead). (D) Quantification. (E) Summary. Scale bars, 5 μm. (F-J) Nkx6 specifies VO motor neuron axon targeting to ventral oblique muscles. (F) Controls have NB7-1 progeny (*NB7-1-gal4 UAS-GFP*) projecting out the ISNd to the ventral oblique muscles. (G) Nkx6 overexpression in the NB7-1 lineage (*NB7-1-gal4 UAS-Nkx6*) have ectopic projections out the ISNd to the ventral oblique muscles. Scale bar: 10 μm. (H) Flipout labeling of single motor neurons in the NB7-1 lineage shows distinct HA+ and V5+ motor neurons in the lineage projecting out the ISNd to the ventral oblique muscles. Scale bar: 7 μm. (I) Nkx6-RNAi in the NB7-1 lineage (*NB7-1-gal4 UAS-Nkx6-RNAi*) lack projections out the ISNd to the ventral oblique muscles. (J) Quantification. (K) Summary.

We next asked whether Nkx6 was necessary and sufficient to generate VO motor neurons that correctly target the ventral oblique muscles. In wild type, NB7-1 always generates a motor neuron that exits the ISNd and targets ventral oblique muscles (Figure 6F; volume of ISNd projections quantified in 6K). Overexpression of Nkx6 in the NB7-1 lineage generates a more robust FasII projection in the ISNd and a corresponding increase in NB7-1 lineage projections into ISNd (Figure 6G; quantified in 6J, S2C). Importantly, the same phenotype is observed when individual neurons within the NB7-1 lineage are uniquely labeled by MCFO, showing that multiple neurons in the lineage are exiting via ISNd (Figure 6H). Conversely, when we use *NB7-1-gal4* to drive expression of *UAS-Nkx6-RNAi*, we detect a loss of ISNd projections in the NB7-1 lineage (Figure 6I; compare to 6F, S2D). We conclude that Nkx6 is necessary and sufficient to generate VO motor neurons in the NB7-1 lineage. Moreover, our data support a linear pathway in which the TTFs Kr/Pdm induce Nkx6 to specify the molecular and morphological characteristics of the VO motor neuron (Figure 6K; see Discussion).

## Discussion

Kr/Pdm co-expression has been detected in several neuroblast lineages (Doe, 2017; Isshiki et al., 2001), but until now there has not been evidence that this TTF combination could specify neuronal identity. We previously showed that the Kr/Pdm window generates a Kr/Pdm GMC (Averbukh et al., 2018), and here we show that this GMC generates an Nkx6+ ventral-projecting motor neuron. How can Kr and Pdm together specify one fate (Nkx6+ VO motor neuron) whereas Kr or Pdm alone specify completely different fates (Eve+ dorsal motor neuron)? It is likely that Kr/Pdm together activate a different suite of target genes than either alone. For example, Kr/Pdm together may directly activate expression of Nkx6, whereas neither alone has that potential. The emergence of single cell transcriptome and ChIP studies (Stuart and Satija, 2019) will help to reveal how the combination of Kr/Pdm TTFs generates different cell fate output compared to Kr or Pdm alone.

The production of Nkx6+ VO motor neuron in Kr/Pdm window interrupts the sequential production of Eve+ dorsal motor neurons in the NB7-1 lineage, resulting in an Eve>Nkx6>Eve alternation of cardinal motor neuron production within the lineage. This is unusual, as in most cases neurons with similar morphology or function are produced together in a lineage. In mammals, progenitors generate neurons first, followed by glia (Kohwi and Doe, 2013); we know of no examples of neuron>glia>neuron production from a single lineage. Similarly, *Drosophila* central brain neuroblast lineages produce the mushroom body γ neurons, then α’/ β’ neurons, and lastly α/β neurons, with no evidence for alternating or interspersed fates (Lee et al., 1999). In the abdominal NB3-3 lineage, the early-born cells are in a mechanosensitive circuit, whereas the late-born cells are in a proprioceptive circuit (Wreden et al., 2017). To our knowledge, the only other example of interleaved cardinal neuronal subtype production is in the *Drosophila* lateral antennal lobe neuroblast lineage, where there is alternation between uniglomerular projection neurons and AMMC projection neurons (Lin et al., 2012). The use of clonal and temporal labeling tools will be needed to examine additional lineages to determine the prevalence of lineages producing temporally interleaved neuronal subtypes as in the NB7-1 lineage.

Overexpression of Kr/Pdm or Nkx6 can induce only two ectopic VO motor neurons within the NB7-1 lineage. Clearly not all neurons in the lineage are competent to respond to these transcription factors. Early-born temporal identities specified by Hb and Kr (U1-U3) are unaffected by Kr/Pdm or Nkx6 overexpression, which is similar to previous data showing that early temporal fates are not affected by overexpression of later TTFs in multiple lineages (Cheesman et al., 2004; Grosskortenhaus et al., 2006; Tran and Doe, 2008). It remains a puzzle why the Kr+ U3 neuron does not switch to a VO fate upon overexpression of Kr/Pdm. There may need to be a equal level of Kr and Pdm to specify VO fate, although this would not explain why Kr/Pdm overexpression converts the Pdm+ U4 motor neuron to a VO fate. Alternatively, there may be an early chromatin landscape that blocks access to relevant Pdm target loci.

Nkx6 and Eve have cross-repressive interactions (Broihier et al., 2004), but with some limitations: early-born Eve+ motor neurons are not affected by Nkx6 overexpression (our work and Broihier et al., 2004; Cheesman et al., 2004). There is even sporadic expression of Nkx6 in the Eve+ U2 motor neuron (data not shown), but in these neurons it has no effect on Eve expression, nor does it promote targeting to ventral oblique muscles. There appears to be a mechanism to block endogenous or overexpressed Nkx6 function in the early lineage of neuroblasts producing Eve+ motor neurons. The mechanism “protecting” early-born Eve+ neurons from Nkx6 repression of Eve is unknown. Early lineages may lack an Nkx6 cofactor; Nkx6 could act indirectly via an intermediate transcription factor missing in early lineages; the early TTFs Hb or Kr may block Nkx6 function; or the *eve* locus could be in a subnuclear domain inaccessible to Nkx6.

Nkx6 promotes motor neuron specification in both *Drosophila* and vertebrates (Broihier et al., 2004; Cheesman et al., 2004; Sander et al., 2000; Sander et al., 2000). In *Drosophila*, loss of Nkx6 reduces ventral projecting motor neuron numbers and increases the number of Eve+ neurons, while overexpression increases ventral projecting motor neuron numbers at the expense of Eve+ neurons (our work and Broihier et al., 2004). In vertebrates the sole Nkx6 family members Nkx6.1/Nkx6.2 appear to play a broader role in motor neuron specification. Nkx6.1/Nkx6.2 show early expression throughout the pMN domain; mice mutant for both Nkx6 family members lack most somatic motor neurons; and Nkx6.1 overexpression in chick or zebrafish can induce ectopic motor neurons (Briscoe et al., 2000; Cheesman et al., 2004; Sander et al., 2000). It would be interesting to investigate whether vertebrate Nkx6.1/Nkx6.2 are required to suppress a specific motor neuron identity, similar to the antagonistic relationship between Nkx6 and Eve in *Drosophila*.

Neuroblasts in all regions of the *Drosophila* CNS (brain, ventral nerve cord, optic lobe) use TTF cascades to generate neuronal diversity (Allan and Thor, 2015; Doe, 2017; Holguera and Desplan, 2018; Miyares and Lee, 2019), yet less is known about TTF target genes. We identified a linear pathway from Kr/Pdm to Nkx6 which specifies VO motor neuron identity, making Nkx6 one of the few TTF target genes acting to link TTFs to neuronal identity. TTFs act transiently to generate long-lasting neuronal identity, and thus TTFs could act by two non-mutually exclusive mechanisms: inducing more stable combinatorial codes of transcription factors that consolidate neuronal identity, or by altering the chromatin landscape to have a heritable, long lasting effect. Our finding that transient Kr/Pdm induces stable expression of Nkx6 to promote VO motor neuron identity supports the former mechanism. Identification of Nkx6 target genes would give a more comprehensive understanding of TTF specification of neuronal identity. In addition, identification of factors acting downstream of Hb, Kr and other TTFs will be an important goal for the future (Sen et al., 2019).

The results presented in this work lead to several interesting directions. Other embryonic VNC lineages exhibit a Kr/Pdm window; does this window generate neurons in these lineages? Are there common features to neurons born in the Kr/Pdm window? Furthermore, do ectopic VO neurons make functional presynapses with the ventral oblique muscles, and do they have the normal inputs to their dendritic postsynapses? In only a few cases has it been shown the TTF-induced neurons are functionally integrated into the appropriate circuits (Meng et al., 2019). Kr and Pdm orthologs have been identified in vertebrates, although their function is still poorly defined. Looking for dual expression of Kr and Pdm orthologs in vertebrates may reveal a role in specifying temporal identity, similar to evidence for Hb and Cas TTFs having vertebrate orthologs that specify temporal identity (Alsiö et al., 2013; Elliott et al., 2008; Mattar et al., 2015; Mattar et al., 2020).

## Methods

### Fly Stocks

Male and female *Drosophila melanogaster* were used. The chromosomes and insertion sites of transgenes (if known) are shown next to genotypes. Previously published gal4 lines, mutants and reporters used were: *NB7-1-gal4*^*KZ*^ (II) (Seroka and Doe, 2019); *en-gal4* (Isshiki et al., 2001; Pearson and Doe, 2003; Schmid et al., 1999); *10XUAS-IVS-myr::sfGFP-THS-10xUAS(FRT.stop)myr::smGdP-HA* (RRID:BDSC_62127); *UAS-nkx6* (RRID:BDSC_9932); *UAS-nkx6*^*RNAi*^ (RRID:BDSC_61188); *UAS-Kr* (II and III) (Cleary and Doe, 2006; Isshiki et al., 2001); *UAS-Pdm2* (II and III) (Grosskortenhaus et al., 2006; Tran and Doe, 2008); *hs-FLPG5.PEST.Opt* (RRID:BDSC_77140); *hs-FLPG5.PEST, 10xUAS(FRT.stop)myr::smGdP-OLLAS 10xUAS(FRT.stop)myr::smGdP-HA 10xUAS(FRT.stop)myr::smGdP-V5-THS-10xUAS(FRT.stop)myr::smGdP-FLAG* (RRID:BDSC_64086).

### Immunostaining and imaging

Primary antibodies were: mouse anti-Eve (5 μg/mL, DSHB, 2B8), rabbit anti-Eve #2472 (1:100, Doe Lab), mouse anti-FasII (1:50, DSHB, 1D4), chicken anti-GFP (1:1000, RRID:AB_2307313, Aves Labs, Davis, CA), rabbit anti-HA epitope tag, DyLight™ 549 conjugated (1:100, Rockland, 600-442-384, Limerick, PA), rat anti-HA (1:100, MilliporeSigma, 11867423001, St. Louis, MO), guinea pig anti-Hey (1:1000, gift from S. Bray, University of Cambridge), guinea pig anti-Kr (1:500, Doe lab), rat anti-Nkx6 (1:500, gift from J. Skeath, Washington University in St. Louis), rat anti-Pdm2 (abcam, ab201325, Cambridge, MA), guinea pig anti-Runt (1:1000, gift from C. Desplan, New York University), chicken anti-V5 (Bethyl Laboratories, Inc. A190-118A, Centennial, CO), rabbit anti-Zfh1 (1:1000, gift from R. Lehman, New York University), guinea pig anti-Zfh1 (1:1000, gift from J. Skeath, Washington University in St. Louis), and. Fluorophores-conjugated secondary antibodies were from Jackson ImmunoResearch (West Grove, PA) and were used at 1:200.

Embryos and the whole newly hatched larvae were fixed and stained as previously described (Seroka and Doe, 2019). In some cases, larval brains were dissected in PBS, fixed in 4% paraformaldehyde, and then stained by following protocols as described (Seroka and Doe, 2019). The samples were mounted either in Vectashield (Vector Laboratories, Burlingame, CA) or DPX (Meissner et al., 2018).

Images were captured with a Zeiss LSM 710 or Zeiss LSM 800 confocal microscope with a *z*-resolution of 0.35 μm. Due to the complex 3-dimensional pattern of each marker assayed, we could not show NB7-1 progeny marker expression in a maximum intensity projection, due to irrelevant neurons in the z-axis obscuring the neurons of interest; thus, NB7-1 progeny were montaged from their unique z-axis position while preserving their X-Y position. This was done in Figures 1, 4, and 6. Images were processed using the open-source software FIJI (https://fiji.sc). Figures were assembled in Adobe Illustrator (Adobe, San Jose, CA). Three dimensional reconstructions, and level adjustments were generated using Imaris (Bitplane, Zurich, Switzerland). Any level adjustment was applied to the entire image.

### Quantification of Normalized ISNd volume

In order to generate a reliable volume metric for determining changes to the size of the ISNd nerve, FasII staining was used to identify SNc in a 3D IMARIS volume of the confocal image stack in order to normalize for variance in embryonic age and development. SNc was chosen to normalize against as we observed it to be unaffected by our genetic manipulations. Default parameters were used to generate a surface over the FasII channel labeling SNc and ISNd respectively, generating precise volume measurements. We then used these volume measurements to generate a ISNd/SNc ratio which was used for statistical analysis (see Supplemental Figure 1). This metric exhibited low variance across the wildtype controls, allowing for accurate comparison to genetically manipulated embryos.

### Statistics

Statistical significance is denoted by asterisks: ****p<0.0001; ***p<0.001; **p<0.01; *p<0.05; n.s., not significant. The following statistical tests were performed: Mann-Whitney U Test (non-normal distribution, non-parametric) (two-tailed p value) (Figures 4C, F, I, J, 6D); Welch’s t-test (normal distribution, non-parametric) (two-tailed p value) (Figures 5C, 6J). All analyses were performed using Prism8 (GraphPad). The results are stated as mean ± s.d., unless otherwise noted.

## Acknowledgements

We thank Alice Schmid for the DiI clone in Figure 2A, Ellie Heckscher, Judith Eisen, and Emily Heckman for comments on the manuscript, Sarah Bray, Claude Desplan, Ruth Lehman, Jim Skeath, and Stefan Thor for fly stocks and/or antibodies. We thank Bloomington Stock Center (NIH P40OD018537) for fly stocks, and DSHB for antibodies. This work was funded by HHMI (S-LL and CQD), NIH R01 HD27056 (CQD), and T32 HD007348 (AQS).

## Author contributions

A.Q.S., S.-L.L and C.Q.D. conceptualized the work. A.Q.S., R.M.Y. and S.-L.L. performed experiments and analyzed results. All authors contributed to writing the manuscript.

## Figure legends

**Figure S1.**
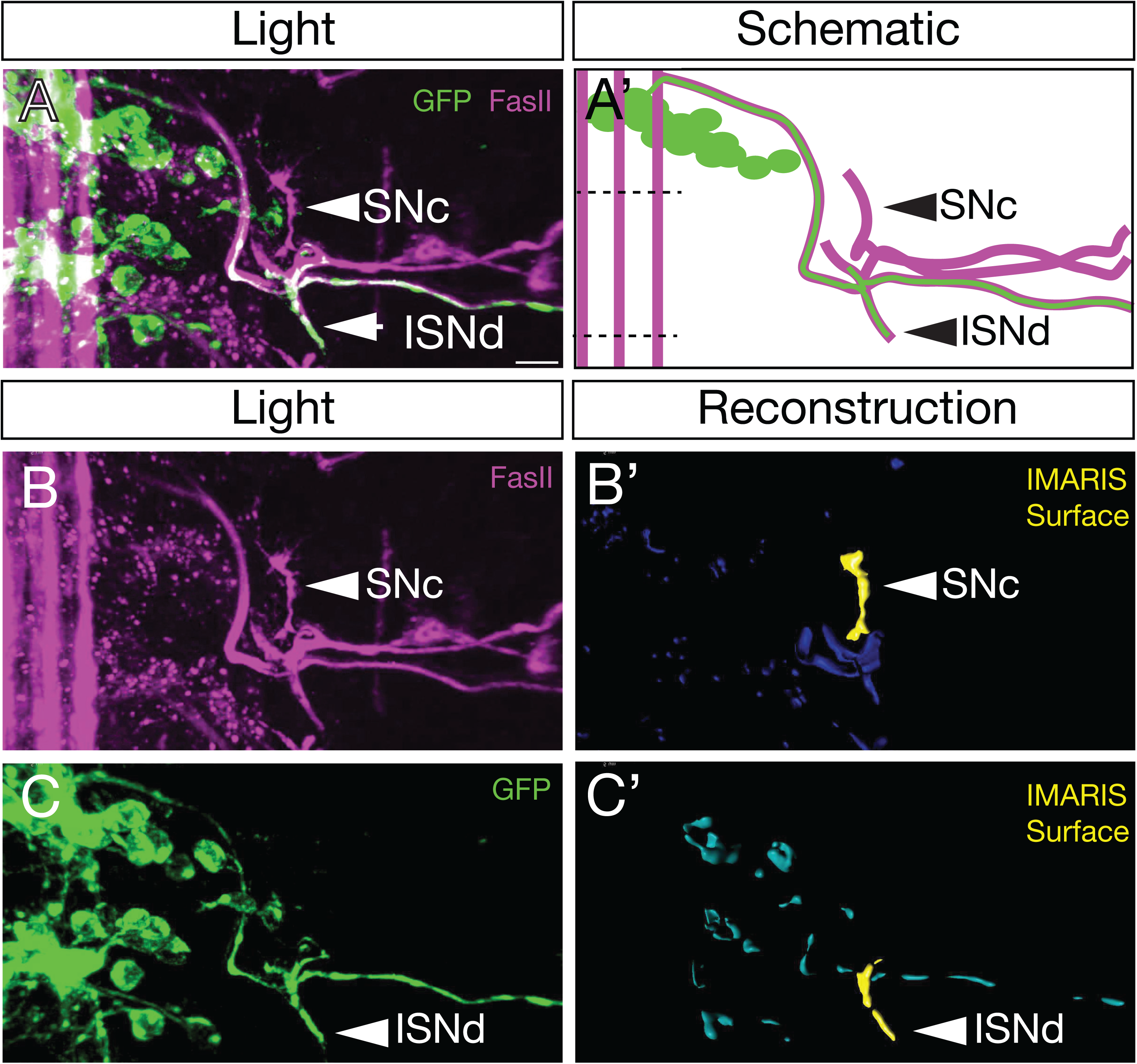
Methodology for quantifying ISNd and SNc motor neuron localization. (A,A’) The volume of the ISNd was normalized to that of SNc to account for slight differences in embryo staging. ISNd and SNc were identified by the pan-motor axon marker FasII (magenta) in embryos expressing GFP (green) in the NB7-1 lineage (*NB7-1-gal4 UAS-GFP*). (B,B’) FasII (magenta) was used to identify SNc in a maximum intensity projection, and the volume quantified using the Imaris Surface function. (C,C’) FasII (magenta) was used to identify ISNd in a maximum intensity projection, and the volume quantified using the Imaris Surface function.

**Figure S2.**
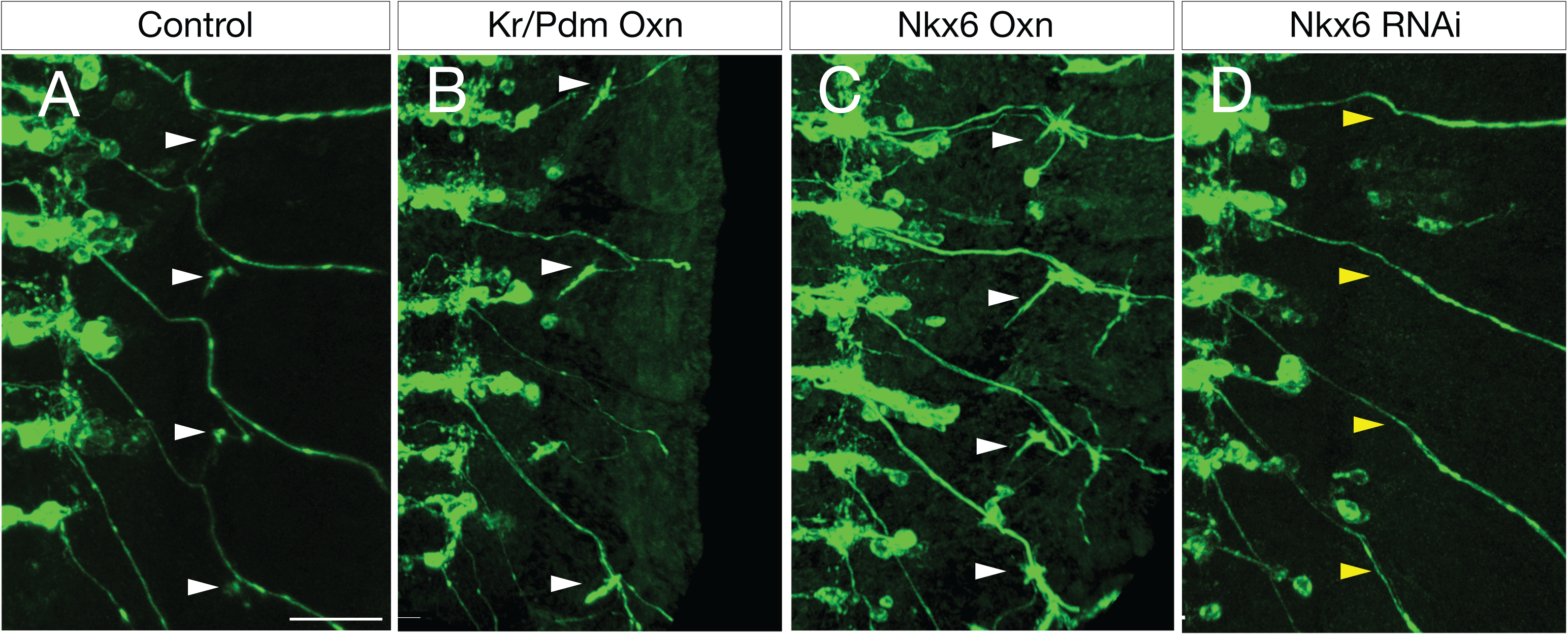
Nkx6 induces ectopic VO motor neurons targeting ventral oblique muscles. (A) Control NB7-1 progeny (*NB7-1-gal4 UAS-GFP*) show precise innervation of the ISNd and ventral oblique muscles (arrowhead). Scale bar: 15 μm. (B) NB 7-1 progeny misexpressing Kr and Pdm (*NB7-1-gal4 UAS-GFP UAS-Kr UAS-Pdm*) demonstrate overexpansion of the ISNd domain and unusual increase in ISNd size in most hemisegments (arrowhead). (C)Overexpression of Nkx6 in the NB7-1 lineage (*NB7-1-gal4 UAS-Nkx6 UAS-GFP*) leads to excessive, broad, and disorganized innervation of ventral oblique muscles (arrowhead). (D) Knockdown of Nkx6 in the NB7-1 lineage (*NB7-1-gal4 UAS-Nkx6-RNAi*) results in loss of ventral motor projections to ISNd (yellow arrowhead).

